# Replication stress links Geminin depletion to centrosome amplification

**DOI:** 10.64898/2026.06.30.735730

**Authors:** Inês B. Santos, David M. Glover

## Abstract

Both DNA replication origins and centrioles undergo licensing events that restrict their duplication to once per cell cycle; yet how defects in DNA replication licensing are coupled to centrosome homeostasis remains poorly understood. Here, we show that depletion of the replication licensing inhibitor Geminin in proliferating mouse embryonic fibroblasts induces centrosome amplification and impairs primary cilium assembly without whole–genome re–replication. Instead, knockdown of Geminin triggers a replication stress–associated DNA damage response that activates the ATR–Chk1– Wee1 checkpoint axis, enabling untimely centriole licensing and centrosome duplication. Ultrastructure expansion microscopy reveals increased distance between disengaged centrioles and centriole pairs following Geminin depletion, which is restored by Plk1 inhibition. Centrosome amplification induced by independent genetic and pharmacological sources of replication stress is similarly suppressed by inhibition of ATR or Plk1, indicating that this mechanism is not restricted to Geminin loss. Together, our findings identify replication stress as a signaling module that couples defective DNA replication licensing to centrosome amplification and reveal a pathway coupling genome surveillance to centrosome duplication.

## Introduction

Accurate duplication of both the genome and centrosome is essential to preserve genomic integrity. Each centrosome consists of two barrel–shaped, orthogonally oriented centrioles, surrounded by a protein–rich pericentriolar matrix (PCM). The centriole duplication cycle is triggered during the telophase of the previous cell cycle by centriole disengagement under the control of mitotic kinase Plk1 (Polo–like kinase 1) and the protease, Separase [1][2][3]. As centrioles disengage, they lose their orthogonal orientation and steric hindrance for procentriole formation, thus licensing them for duplication during the next S phase. The duplicated centrosomes, each containing a pair of engaged centrioles, separate at mitotic onset to orient the bipolar mitotic spindle.

DNA replication is tightly regulated to occur once per cell cycle. This is achieved in two steps. First, in late mitosis/early G1, replication origins are licensed through assembly of a pre–replication complex (preRC). The preRC assembles sequentially through binding of the origin recognition complex (Orc) to DNA origins, followed by Cdc6 recruitment, which then recruits the replicative helicase, Mcm2–7, and the licensing factor Cdt1 [4]. The second step occurs during S phase, when replication forks fire at licensed origins. Multiple mechanisms exist to control Cdt1 protein levels: ubiquitin mediated–degradation of Cdt1 blocks DNA licensing in S phase; phosphorylation of Cdt1 by Cyclin–dependent kinase 1 (Cdk1) blocks Mcm2–7 recruitment in G2 phase; and the binding of Cdt1 to the inhibitory protein Geminin in S and G2 phases blocks DNA licensing in metazoans [5][6].

In their initial identification of Geminin, McGarry and Kirschner mapped a 9 amino–acid (aa) destruction box consensus in its N–terminus that is recognized by mitotic anaphase–promoting complex (APC) to control Geminin levels. At the G1/S transition, the APC is inactivated, allowing Geminin to accumulate, preventing licensing of new origins. Because Geminin has no effect on elongation, DNA replication continues to completion. At the metaphase–anaphase transition, the APC is activated, mediating ubiquitination and the abrupt degradation of Geminin, thereby permitting another round of replication to initiate in the following cell cycle [7].

These processes are coordinated through cell–cycle dependent mechanisms that prevent re–licensing of DNA origins or centrioles. Failure to maintain such mechanisms results in replication stress, genomic instability and centrosome amplification, common features of human cancers and developmental disorders [8][9]. Indeed, inherited mutations in Geminin and other factors associated with the initial steps of DNA replication fork assembly result in Meier–Gorlin Syndrome (MGS), a rare genetic disorder conferring microcephaly and growth defects [10][11].

Loss of Geminin has been reported to induce DNA re–replication and centrosome amplification in multiple experimental systems, yet the mechanistic relationship between these phenotypes remains unclear [12]. In particular, it has not been resolved how centrosome amplification arises under such conditions. Here, we show that depletion of Geminin in proliferating mouse embryonic fibroblasts (MEFs) induces robust centrosome amplification without whole–genome duplication. Instead, Geminin loss triggers a replication stress state that activates a checkpoint response dependent on ATR (Ataxia–Telangiectasia Mutated and RAD3–related) that promotes centriole amplification through a Wee1–Plk1 dependent pathway. We further show that independent genetic and pharmacological induction of replication stress similarly promotes ATR– and Plk1–dependent centrosome amplification. Our findings identify replication stress as a signal coupling defective DNA licensing to centriole amplification and provide a mechanistic framework linking genome surveillance pathways to centrosome duplication mechanisms.

## Results and discussion

To investigate the function of Geminin in cell cycle progression, we examined the effects of inhibiting its expression using short interfering RNA (siRNA) in MEFs. Centrosome number was assessed by co–staining to reveal the pan–centriolar marker Cep152 and **γ**–tubulin to count the total number of centriole pairs, or by co–staining to reveal the distal appendage protein Cep164 and **γ**–tubulin to quantify mother centrioles. Control MEFs predominantly contained one or two centrosomes, reflecting normal cell cycle progression. In contrast, Geminin–depleted cells exhibited increased numbers of centrosomes that were positive for Cep152 (2.14±1.20% in control vs. 23.09±2.59% in Geminin–depleted cells, Figure 1A/C), and an increase in cells that had greater than two Cep164 positive mother centrioles (0.37±0.22% in control vs. 10.82±2.98% in Geminin–depleted cells, Figure 1B/D). Efficient depletion of Geminin was confirmed both at the transcript and at the protein level, and four independent siRNA oligonucleotides produced comparable centrosome amplification phenotypes, demonstrating that the observed defects were not attributable to off–target effects (Supplemental Figure 1).

**Figure 1.**
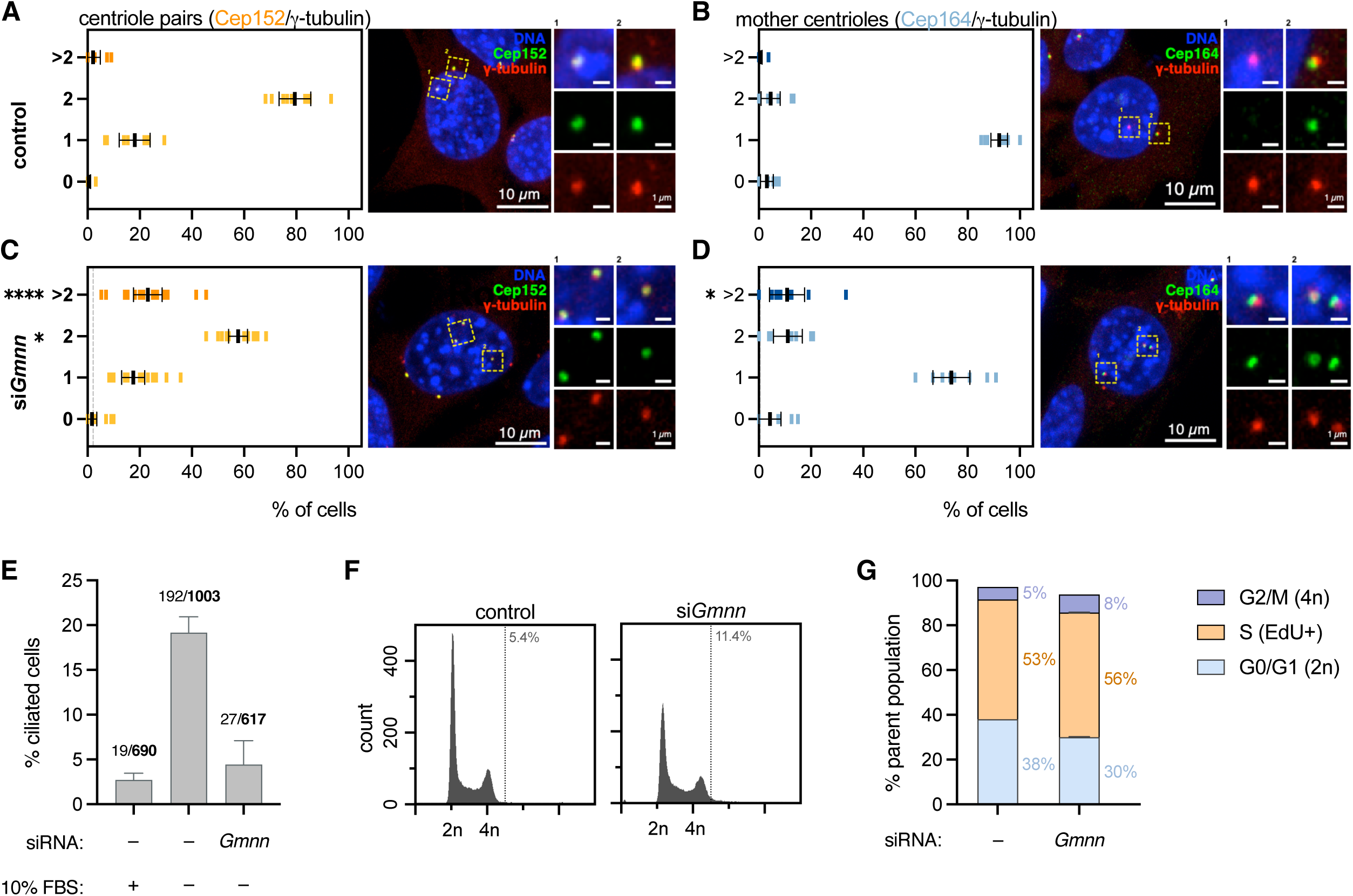
Geminin depletion induces centrosome amplification and loss of primary cilia in proliferating mouse embryonic fibroblasts. (A) Graphic representation of the percentage of cells showing 0, 1, 2 and more than 2 (>2) centriole pairs, per cell. Mouse embryonic fibroblasts (MEFs) were stained to reveal Cep152 (green), γ–tubulin (red), *a*–tubulin (not shown) and DNA (blue). Colocalization of Cep152 with γ–tubulin identifies centriole pairs/centrosomes. (B) Graphic representation of the percentage of cells showing 0, 1, 2 and more than 2 (>2) mother centrioles, per cell. MEFs are stained to reveal Cep164 (green), γ–tubulin (red), *a*–tubulin (not shown) and DNA (blue). Colocalization of Cep164 with γ–tubulin identifies mother centrioles. (C) As in (A). MEFs were subjected to Geminin siRNA, and the number of centriole pairs/centrosomes was analyzed 3 days post–transfection (dpt). (D) As in (B). MEFs were subjected to Geminin siRNA, and the number of mother centrioles was analyzed 3 dpt. (E) Control or Geminin–depleted MEFs were serum starved for 24 hours and processed to analyze primary cilia assembly using anti–acetylated tubulin and anti–Arl13B antibodies. The percentage of ciliated cells was measured for each group. Number of ciliated cells/total numbers of cells analyzed are depicted above each bar. (F) Control or Geminin– depleted MEFs were analyzed by FACS at 3 dpt. FACS profile from a representative experiment and the percentage of cells with DNA content above 4n are shown. (G) FACS profiles described in (F) were used to quantify the percentage of cells in each cell cycle phase. EdU incorporation was used to determine cells actively replicating DNA (S phase), and DNA counts were used to identify 2n (G0/G1 cells) and 4n (G2/M cells) peaks. The distribution of cells in each cell cycle phase is depicted as a percentage of the parental population next to each bar. Data information: In (A–D) the number of centrioles was quantified for each category, and percentage values are shown as vertical bars. Experiments were performed at least 3 times independently. Error bars correspond to mean ± 95% confidence interval (CI). The statistical significance was calculated using the Kruskal–Wallis test, followed by Dunn’s multiple comparisons test for multiple pairwise comparisons. P–values are represented with one (P<0.05), or four (P<0.0001) asterisks. Scale bar, 10 µm; inset scale bar, 1 µm. In (E and G) the average and standard deviation (SD) of three independent experiments are plotted.

Because centrioles also template basal body formation necessary for primary cilium assembly, we next examined whether Geminin depletion compromised this centrosome associated function. Following serum starvation, Geminin–depleted MEFs exhibited a pronounced decrease in primary cilia assembly compared with serum– starved control cells (4.43±1.53% in Geminin–depleted vs. 19.17±1.01% in control cells, Figure 1E), indicating that centrosome amplification was accompanied by an impaired ability of mother centrioles to generate competent basal bodies.

Geminin depletion has previously been proposed to induce centrosome amplification as a consequence of DNA re–replication [12]. To determine if the observed phenotype reflected whole–genome duplication, we analyzed DNA content and thereby, cell cycle distribution by flow cytometry. Geminin depletion resulted in a small accumulation of cells in S/G2, consistent with impaired DNA replication, but only a modest increase in cells with DNA content above 4n (11.4% vs. 5.4%, Figure 1F/G). Importantly, no evidence of a distinct polyploid population was observed, merely a widening of the 4n tail. These observations suggest that in MEFs, Geminin depletion perturbs DNA replication without whole–genome re–replication, raising the possibility that replication stress, rather than polyploidization or cytokinesis failure, underlies centrosome amplification.

To distinguish between the above possibilities, we examined activation of the DNA damage response during S phase by pulse–labelling cells with EdU (5–ethynyl–2′– deoxyuridine) and quantifying markers of DNA damage signaling. We found that Geminin–depleted MEFs exhibited an increase in **γ**–H2AX foci relative to control cells, indicating accumulation of DNA lesions during DNA replication (Figure 2A/B). This response was accompanied by an increase in phosphorylated Chk1 foci, in line with activation of the ATR–dependent replication stress checkpoint (Figure 2A/C).

**Figure 2.**
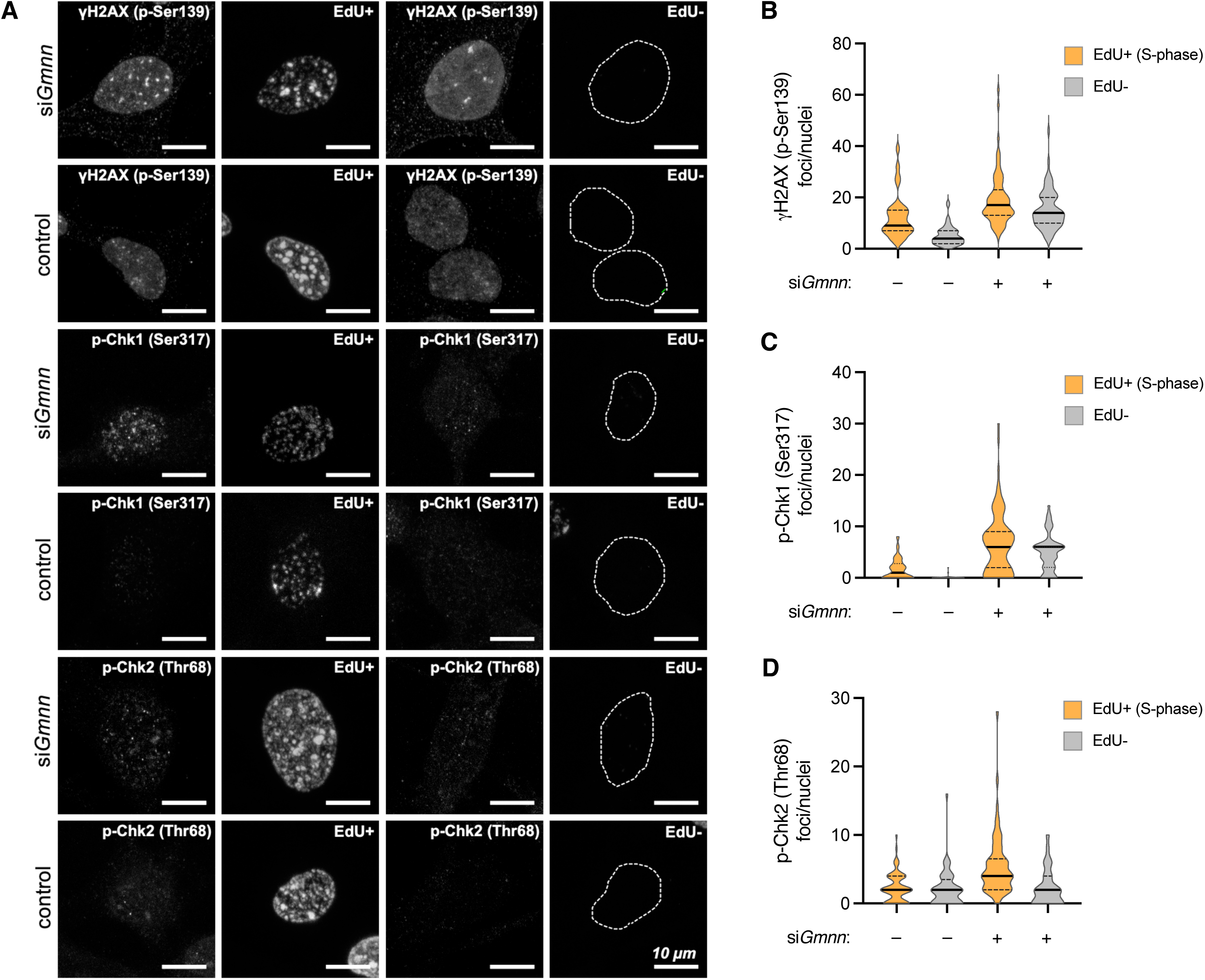
Geminin depletion induces a replication–stress associated DNA damage response without whole genome re–replication. (A) Control and Geminin–depleted MEFs were treated with pulse–labelled EdU for 2 hours prior to fixation, to identify cells actively replicating DNA. Click–it reaction and immunostaining against **γ**–H2AX (phospho–Ser139) or Chk1 (phospho–Ser317) or Chk2 (phospho–Thr68) was carried out as described in the Materials and Methods. Representative images for each antibody are shown. Scale bar, 10 µm. (B) The number of **γ**–H2AX (phospho–Ser139) foci per nuclei was quantified using FIJI software and plotted. (C) The number of Chk1 (phospho–Ser317) foci per nuclei was quantified using FIJI software and plotted. (D) The number of Chk2 (phospho–Thr68) foci per nuclei was quantified using FIJI software and plotted. Data information: Representative images are shown in (A). In (B–D) at least 200 cells per condition were quantified in 3 independent experiments. For each data set, black line represents median and dashed lines represent upper and lower quartiles.

Interestingly, increased numbers of both **γ**–H2AX and phosphorylated Chk1 nuclear foci were present in cells that did not show EdU incorporation, supporting the notion that Geminin–depleted MEFs are in a replication stress state and that DNA damage signaling is not restricted to cells undergoing active DNA synthesis. In contrast, phosphorylation of Chk2 remained largely unchanged following Geminin depletion, with a slight increase in nuclear foci detected in S phase cells (Figure 2A/D). This is likely caused by stalled replication forks progressing to DNA double strand breaks (DSB) and activating the DNA damage response kinase ATM (Ataxia–Telangiectasia Mutated) and its downstream effector Chk2. Taken together, these findings indicate that Geminin depletion preferentially activates an ATR–mediated replication stress response rather than a DNA double–strand break signaling pathway. Activation of the ATR/Chk1 axis is known to suppress the expression of a mitotic gene network during DNA replication to prevent mitotic entry with under–replicated DNA [13][14]. Together with the cell cycle analysis, these observations argue against extensive genome re– replication as the primary consequence of Geminin knockdown in MEFs. Instead, our findings support a model in which deregulated origin licensing generates replication stress, leading to activation of ATR–Chk1 signaling before the onset of centrosome amplification.

We next asked whether activation of the ATR–Chk1 checkpoint contributes functionally to centrosome amplification. To this end, we examined the requirement of DNA damage response kinases for the centrosome amplification process. Geminin–depleted MEFs were treated with selective inhibitors of DNA–PK (DNA–dependent protein kinase), ATM or ATR and centriole numbers were quantified. Inhibition of ATR markedly suppressed Geminin depletion–induced centrosome amplification (ATR_i_: 5.36±1.15% vs. DMSO: 23.09±2.59%, Figure 3A/D). Neither ATM inhibition, nor DNA–PK inhibition, had significant effects upon centrosome amplification following Geminin depletion (ATM_i_: 14.64±1.90%; DNA–PK_i_: 16.23±2.01% vs. DMSO: 23.09±2.59%, Figure 3A–C).

**Figure 3.**
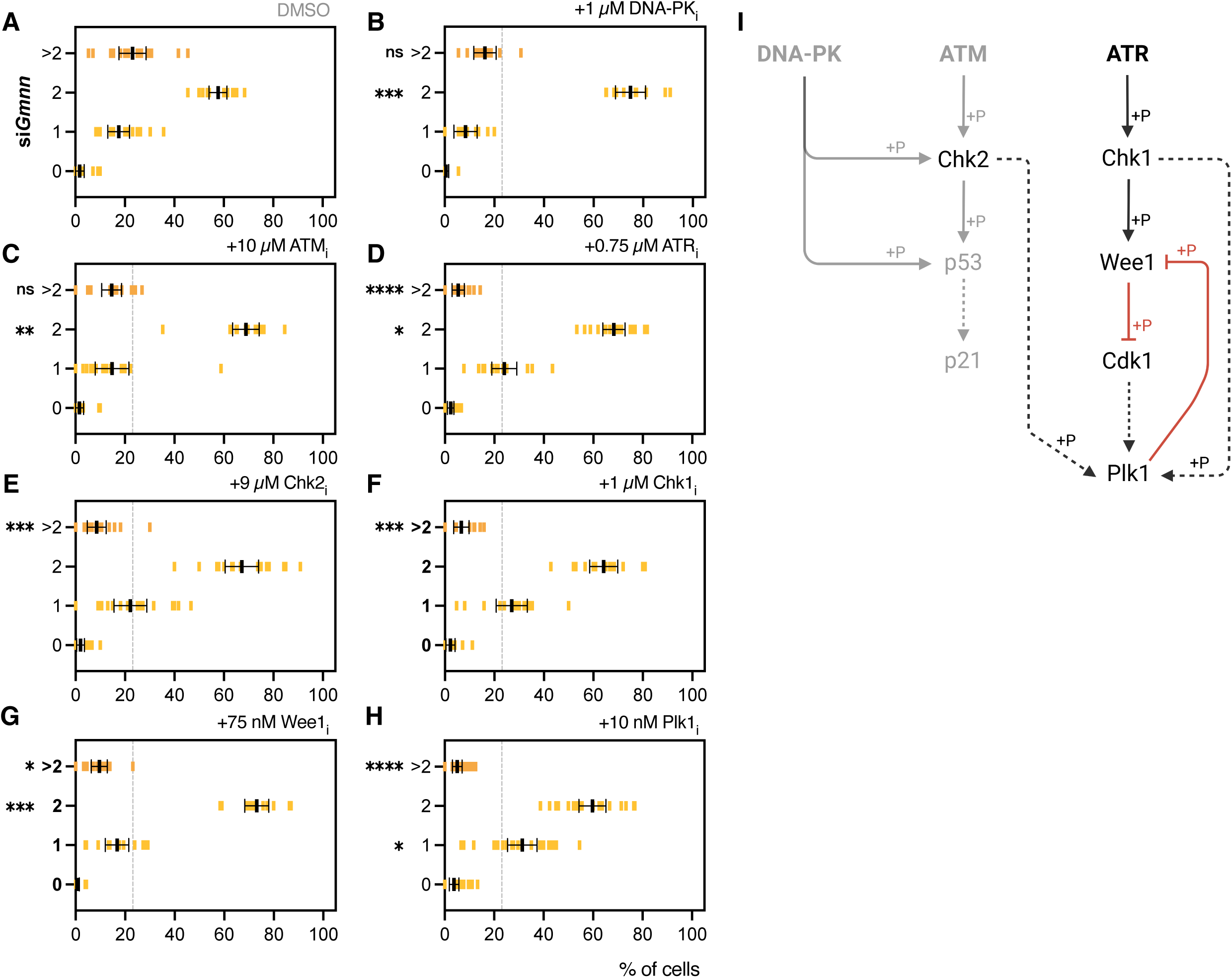
ATR–dependent checkpoint kinase signaling links Geminin depletion– induced replication stress to centriole amplification through Wee1 and Plk1. (A–H) Graphic representation of the percentage of cells showing 0, 1, 2 and more than 2 (>2) centriole pairs/centrosomes, per cell. Geminin–depleted MEFs were stained to reveal Cep152, γ–tubulin, *a*–tubulin and DNA. Colocalization of Cep152 with γ–tubulin identifies centriole pairs/centrosomes. Geminin–depleted MEFs were individually treated with (A) vehicle control, or with inhibitors against the DNA damage response kinases (B) DNA–PK, (C) ATM or (D) ATR, inhibitors of the checkpoint kinases (E) Chk2 or (F) Chk1, and inhibitors against mitotic kinases (G) Wee1 or (H) Plk1. (I) Schematic representation of the DNA damage response molecular pathway regulating premature centriole disengagement upon replication stress. Created in BioRender. Santos, IB. (2026) https://BioRender.com/9nwo5s7. Data information: In (A–H) MEFs were transfected with siRNA against Geminin and centriole number was analyzed at 3 dpt. The number of centrioles was quantified for each category, and percentage values are shown as vertical bars. Experiments were performed at least 3 times independently. Error bars correspond to mean ± 95% confidence interval (CI). The statistical significance was calculated using the Kruskal– Wallis test, followed by Dunn’s multiple comparisons test for multiple pairwise comparisons. P–values are represented with one (P<0.05), two (P<0.01), three (P<0.001) or four (P<0.0001) asterisks. Non–statistical significance is labeled ns. Inhibitors were added to cells in culture, at 1 dpt (inhibitor treatment time: 2 days prior to fixation).

These findings indicate that activation of ATR, rather than the canonical DSB response, is specifically required for centriole amplification following Geminin knockdown. To initiate DNA repair and prevent mitotic entry in cells having damaged DNA, ATR activates the checkpoint kinase Chk1, while ATM activates Chk2. We next examined the contribution of these downstream effector kinases. Pharmacological inhibition of Chk1 suppressed centrosome amplification in Geminin–depleted cells (Chk1_i_: 6.62±1.44% vs. DMSO: 23.09±2.59%, Figure 3A/F), whereas inhibition of Chk2 had a smaller but significant effect in reducing centrosome number (Chk2_i_: 8.54±1.79% vs. DMSO: 23.09±2.59%, Figure 3A/E). The suppression of centrosome amplification by Chk1 inhibition is in line with a requirement for ATR–dependent DNA damage signaling. Interestingly, knockdown of Geminin in *Drosophila* cells has previously been reported to induce re–replication, that can be rescued by silencing of Chk1 but not Chk2 [15][16]. Together, these findings demonstrate that checkpoint signaling not only marks replication stress but is essential to mediate the centrosome amplification response in MEFs. The fact that inhibition of Chk2 but not ATM also had an effect in rescuing Geminin–induced centrosome amplification was surprising, but perhaps not completely unexpected. While no functional relationship has been described until now, Mihaylov *et* al. reported to have isolated Geminin during the analysis of Chk2–associated proteins, in HeLa cells [15]. However, the convergence of checkpoint kinase signaling via Chk2 will require further studies to identify how it becomes activated.

ATR–Chk1 signaling coordinates multiple cell–cycle regulators that control entry into mitosis, including the activation of Wee1 kinase that phosphorylates Cdk1 to prevent mitotic entry and prolong G2 [17]. Accordingly, we found that pharmacological inhibition of Wee1 suppressed the accumulation of supernumerary centrioles in Geminin–depleted MEFs (Wee1_i_: 9.54±1.49% vs. DMSO: 23.09±2.59%, Figure 3A/G), demonstrating its requirement in the ATR checkpoint pathway to inhibit Cdk1. Because Plk1 is a key regulator of centriole disengagement to license centriole duplication, we asked whether its activity was required to mediate centrosome amplification following Geminin depletion. We found that inhibition of Plk1 markedly suppressed the accumulation of supernumerary centrioles in Geminin–depleted MEFs (Plk1_i_: 5.08±0.94% vs. DMSO: 23.09±2.59%, Figure 3A/H). Together, these findings identify a Wee1–Plk1 signaling module essential for the regulation of centrosome homeostasis downstream of replication stress checkpoint activation (Figure 3I).

Having established that centrosome amplification following Geminin depletion depends on replication stress–associated checkpoint signaling, we next asked whether this phenotype was specific to Geminin depletion or represented a broader consequence of replication stress. To address this, we examined centrosome number following two independent replication stress conditions. Previous work demonstrated that mild replication stress induced by nanomolar doses of aphidicolin promotes premature centriole disengagement through ATR/Chk1–dependent modulation of Plk1 activity [18], consistent with studies placing Chk1 upstream of and directly phosphorylating Plk1 [19]. Under these conditions, Plk1 activity is dampened to levels insufficient to promote mitotic entry but sufficient to support premature centriole disengagement via Separase–mediated cleavage of Pericentrin. Inhibition of ATR restores Plk1 activity, thus preventing centriole disengagement [18]. Since premature centriole disengagement licenses centrioles for duplication independently of the normal cell cycle program and is a known mechanism underlying centrosome amplification [2], we asked whether mild replication stress conditions were sufficient to promote centrosome amplification, rather than centriole disengagement alone. We found that low dose aphidicolin treatment resulted in a significant increase in cells with supernumerary centrosomes compared with vehicle–treated controls (aph: 22.51±2.22% vs. DMSO: 2.142±1.20%, Figure 4A–B), which was suppressed by ATR or Plk1 inhibition (aph: 22.51±2.22% vs. aph/ATR_i_: 9.57±1.41%; aph/Plk1_i_: 9.94±1.61%, Figure 4B–D).

**Figure 4.**
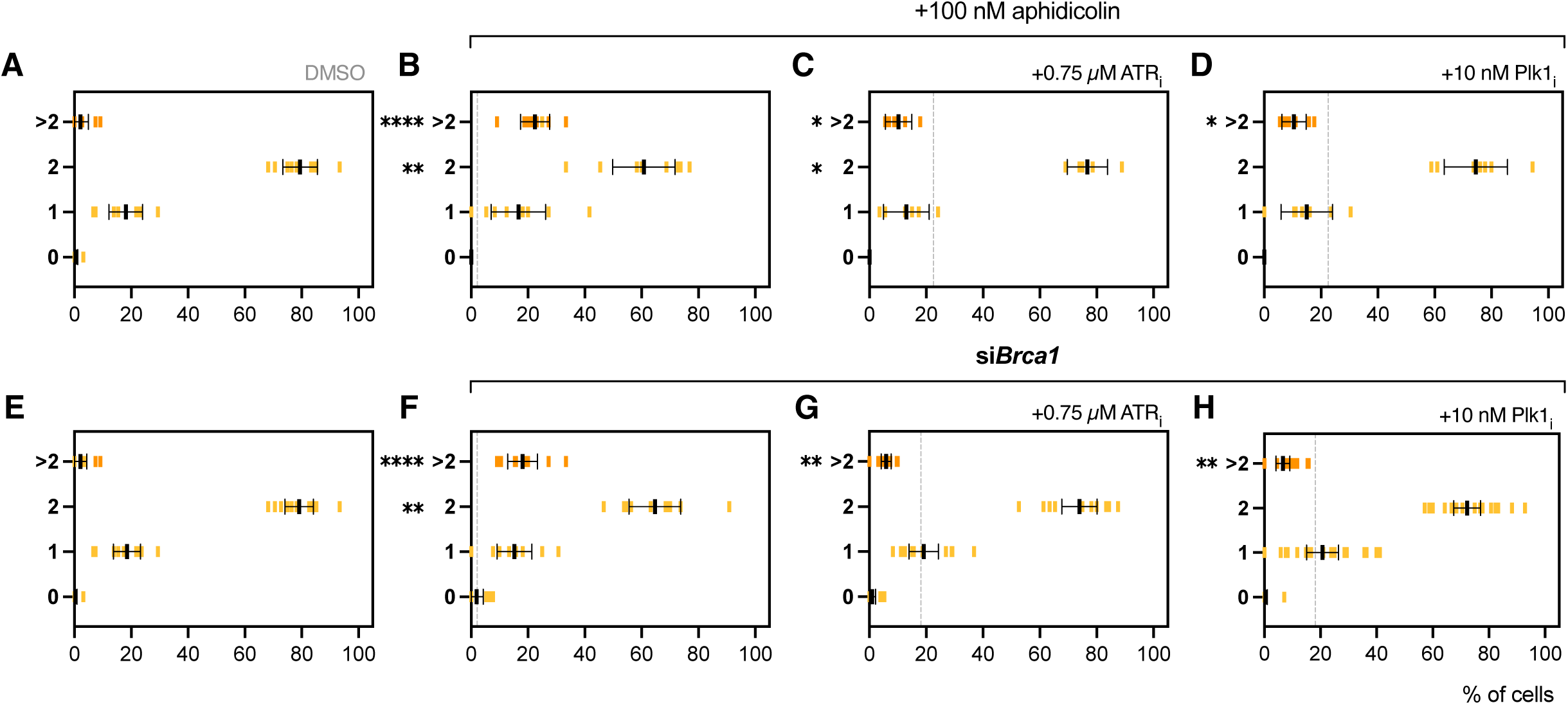
Replication stress-induced centrosome amplification requires ATR and Plk1. (A-H) Graphic representation of the percentage of cells showing 0, 1, 2 and more than 2 (>2) centriole pairs/centrosomes, per cell. Cells were stained to reveal Cep152, **γ**–tubulin, *a*–tubulin and DNA. Colocalization of Cep152 with **γ**–tubulin identifies centriole pairs/centrosomes. MEFs were individually treated with (A) vehicle control, or with inhibitors against (C) ATR, and (D) Plk1, following aphidicolin treatment (B-D). Brca1–depleted MEFs were individually treated with (F) vehicle control, or with inhibitors against (G) ATR, and (H) Plk1. Data information: In (A-D) MEFs were treated with 100 nM aphidicolin. In (F-H) MEFs were transfected with siRNA against Brca1. In (A-H) centriole number was analyzed at 3 dpt. The number of centrioles was quantified for each category, and percentage values are shown as vertical bars. Experiments were performed at least 2 times independently. Error bars correspond to mean ± 95% confidence interval (CI). The statistical significance was calculated using the Kruskal–Wallis test, followed by Dunn’s multiple comparisons test for multiple pairwise comparisons. P–values are represented with one (P<0.05), two (P<0.01), or four (P<0.0001) asterisks. Non–statistical significance is labeled ns. Inhibitors were added to cells in culture, at 1 dpt (inhibitor treatment time: 2 days prior to fixation).

We next asked whether ATR– and Plk1–dependent centrosome amplification could also be observed following an independent genetic perturbation associated with replication stress. To this end, we depleted Brca1, a tumor suppressor with well– established roles in maintaining replication fork stability and whose loss has previously been associated with centrosome amplification [20][21]. Brca1–depleted cells exhibited an increased proportion of cells with supernumerary centrosomes compared with control cells (2.08±1.00% in control vs. 18.13±2.32% in Brca1–depleted cells, Figure 4E–F). Importantly, centrosome amplification following Brca1 depletion was suppressed by inhibition of either ATR or Plk1 (ATR_i_: 5.95±0.80%; Plk1_i_: 6.65±1.16% vs. DMSO: 18.13±2.32%, Figure 4F–H). Together these findings show that ATR– and Plk1–dependent centrosome amplification is not restricted to Geminin depletion but is also observed following independent genetic and pharmacological perturbations associated with replication stress.

Having established that ATR and Plk1 are required for centrosome amplification following distinct perturbations of replication stress, we next sought to determine how such signaling affects centriole organization. To address this, we examined centriole architecture using expansion microscopy. The increased frequency of disengaged centrioles we observed in Geminin–depleted cells suggests that disengagement is premature compared with control cells. Geminin depletion resulted in a two–fold increase in the distance between disengaged centrioles (si*Gmnn*: 4.30±0.53 µm vs. control: 2.06±0.35 µm, Figure 5A/D) and a four–fold increase in the distance between centriole pairs (si*Gmnn*: 12.37±1.42 µm vs. control: 3.00±0.54 µm, Figure 5B/D) compared with control cells. Both phenotypes were suppressed by Plk1 inhibition, with inter–pair distances returning toward control levels, demonstrating that the altered spatial organization of centrioles following Geminin depletion is Plk1–dependent. We also examined engagement within individual centriole pairs by measuring the distance between each centriole and its associated procentriole. Engaged centriole pairs from Geminin–depleted cells show a trend toward increased centriole–procentriole distance (si*Gmnn*: 0.31±0.06 µm vs. control: 0.20±0.02 µm, Figure 5C/D). Together, these measurements reveal a Plk1–dependent increase in centriole separation following Geminin depletion and support premature centriole disengagement as a mechanism contributing to untimely centriole licensing preceding centrosome amplification.

**Figure 5.**
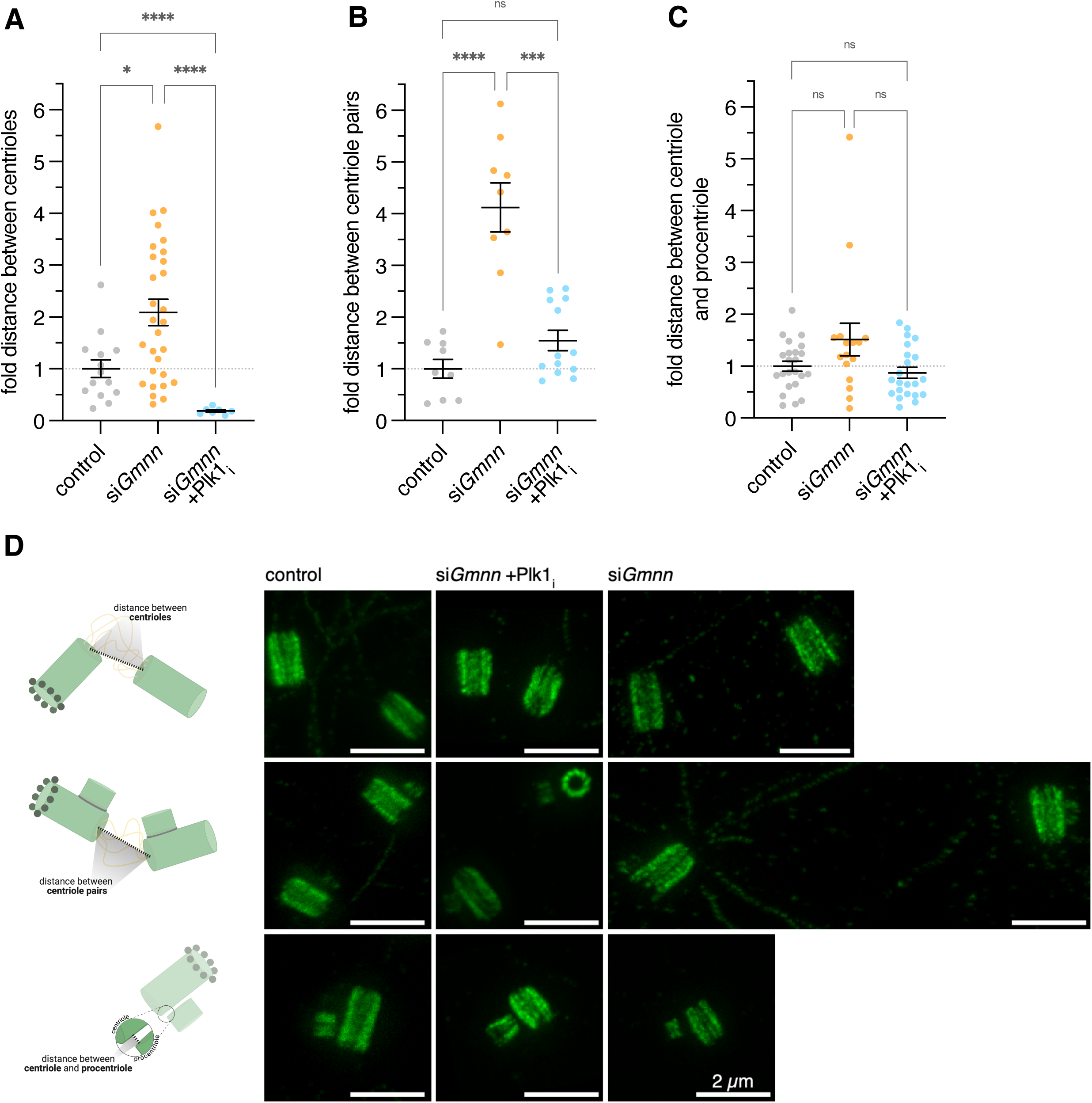
Geminin depletion promotes Plk1–dependent centriole separation. (A–C) Graphic representation of the fold distance measurements in control, Geminin– depleted (si*Gmnn*) and Geminin–depleted MEFs treated with Plk1 inhibitor (si*Gmnn* +Plk1_i_). The distance between (A) disengaged centrioles, (B) centriole pairs and (C) centriole–procentriole was quantified using FIJI software and plotted. (D) MEFs were stained to reveal acetylated–tubulin (green), Cep164 (distal end marker, not shown) or rootletin (proximal end marker, not shown) and DNA (not shown). Representative ultrastructure expansion microscopy (U–ExM) images and schematic representation of the measurement quantified in (A–C). Created in BioRender. Santos, IB. (2026) https://BioRender.com/vo0kjxw. Data information: In (A–D) MEFs were transfected with siRNA against Geminin and cells were fixed at 3 dpt. Plk1 inhibitor was added to cells in culture at 1 dpt (inhibitor treatment time: 2 days prior to fixation). Centriole structure was analyzed by U–ExM. Experiments were performed at least 3 times independently. In (A–C) error bars correspond to mean ± standard error of mean (SEM). Statistical analyses were performed as described in Methods. P–values are represented with one (P<0.05), three (P<0.001) or four (P<0.0001) asterisks. Non–statistical significance is labeled *ns*. In (D) scale bar, 2 µm.

Together, our findings support a model in which loss of Geminin leads to defective DNA replication licensing and induces replication stress that activates ATR/Chk1 checkpoint signaling, that we hypothesize may lead to Wee1/Plk1–dependent premature centriole disengagement and subsequent centrosome amplification (Figure 3I).

Faithful duplication of the genome and centrosome depends on the coordinated licensing of both DNA replication origins and centrioles. Although Geminin has long been recognized as a key inhibitor of DNA origin licensing, how its loss promotes centrosome amplification has been unresolved. Our study demonstrates that, in proliferating MEFs, Geminin depletion induces centrosome amplification not through extensive genome re–replication, but through activation of a replication stress response. We identify ATR–dependent checkpoint signaling as the molecular link between defective DNA replication licensing and centriole amplification and show that this pathway requires downstream regulation of the Wee1/Plk1 axis to promote premature centriole disengagement.

Centrosome amplification following Geminin depletion has generally been interpreted as a secondary consequence of DNA re–replication or polyploidization [12]. While such outcomes have been reported in transformed human cell lines and other experimental systems, our results indicate that this is not the predominant response in embryonic fibroblasts. Instead, Geminin depletion results in a modest accumulation of cells in S/G2 together with activation of **γ**–H2AX and phospho–Chk1, but without the emergence of a substantial polyploid population. This distinction has important implications because it uncouples centrosome amplification from changes in DNA content and instead implicates replication stress–associated checkpoint signaling as the initiating event. Thus, the checkpoint response to Geminin loss may generate distinct outcomes depending on cellular context and/or cell type: facilitating re– replication under conditions permissive for origin re–firing in transformed cell lines, while promoting centrosome amplification through centriole licensing in embryonic fibroblasts.

Our data demonstrates that ATR and Chk1 are required for Geminin depletion–induced centrosome amplification, establishing that replication stress signaling, rather than being concurrent, actively drives this phenotype. Previous studies have reported physical association between Geminin and Chk2, raising the possibility that crosstalk between checkpoint pathways contributes to centrosome homeostasis [15]. Whether this reflects convergence between ATR– and ATM–dependent signaling or an independent role for Chk2 in centrosome biology remains to be determined.

Importantly, the relationship between replication stress signaling and centrosome amplification is not restricted to Geminin depletion. Independent pharmacological and genetic perturbations of DNA replication are able to similarly induce centrosome amplification, extending this response beyond Geminin loss. Thus, distinct perturbations of DNA replication, including deregulated origin licensing, replication fork stalling and replication fork instability, can promote centrosome amplification with a shared requirement for ATR and Plk1 activity.

Mechanistically, our data position a Wee1–Plk1 module downstream of ATR–Chk1 in regulating centriole licensing. At mitotic exit, centriole disengagement licenses centrioles for duplication in the subsequent cell cycle [2]. Plk1 is well established as the regulator of this process, as well as of centrosome disjunction later in the cell cycle [22]. Yet recent work has suggested that replication stress can also modulate Plk1 activity to induce premature centriole disengagement during G2 [18]. Our observations are in accord with this model. U–ExM revealed increased separation between disengaged and centriole pairs under Geminin–depleted conditions, which was restored by Plk1 inhibition. Together, these findings support a model in which replication stress signaling lowers the threshold for the licensing of centrioles, allowing disengaged centrioles to re–duplicate independently of normal cell–cycle progression.

Replication stress is increasingly recognized as a defining feature of oncogene activation, of inherited defects in DNA replication, and of numerous developmental disorders [23][24]. Centrosome amplification is likewise common in these conditions, although the mechanistic relationship between the two has remained poorly understood. Our findings suggest that centrosome amplification need not arise as a passive consequence of genome instability but may instead represent an active cellular response to replication stress mediated by checkpoint signaling.

In summary, we propose a model in which loss of Geminin leads to defective DNA replication licensing that activates an ATR/Chk1–dependent replication stress response that engages Wee1– and Plk1–dependent cell cycle regulation to promote premature centriole disengagement and thereby, centrosome amplification. More broadly, our study identifies replication stress signaling as a mechanism linking perturbations of DNA replication to centrosome duplication, thereby providing a conceptual framework linking genome surveillance pathways with centrosome homeostasis.

## Material and Methods

### Cell culture and transfection

Mouse embryonic fibroblasts (MEFs) were prepared as described previously [25], and cultured in Dulbecco’s Modified Eagle Medium (DMEM) supplemented with 10% fetal bovine serum (FBS). To knock down the expression of each gene, reverse transfection of siRNA into MEFs was performed in a 24–well dish. The transfection mixture comprised 6 pmol siRNA duplex and 1 µl Lipofectamine RNAiMAX (Invitrogen #13778075) in 100 µl Opti–MEM (Gibco #31985070), per well. Liposomes were gently mixed, added to each well, and incubated for 20 minutes at room temperature. MEFs were resuspended to 0.5x10^5^ cells/mL in DMEM, and 500 µl of cell suspension was added per well and incubated with the transfection mix. siRNA used are listed in *Table 1*. Inhibitors (or DMSO control) were added with 600 µl of fresh culture media 1 day post–transfection (dpt). Inhibitors and concentrations are listed in *Table 2*. Cell proliferation, centriole analysis and cell–cycle distribution were assayed at 3 dpt. DNA damage accumulation was assayed 36 hours post–transfection.

**Table 1.**
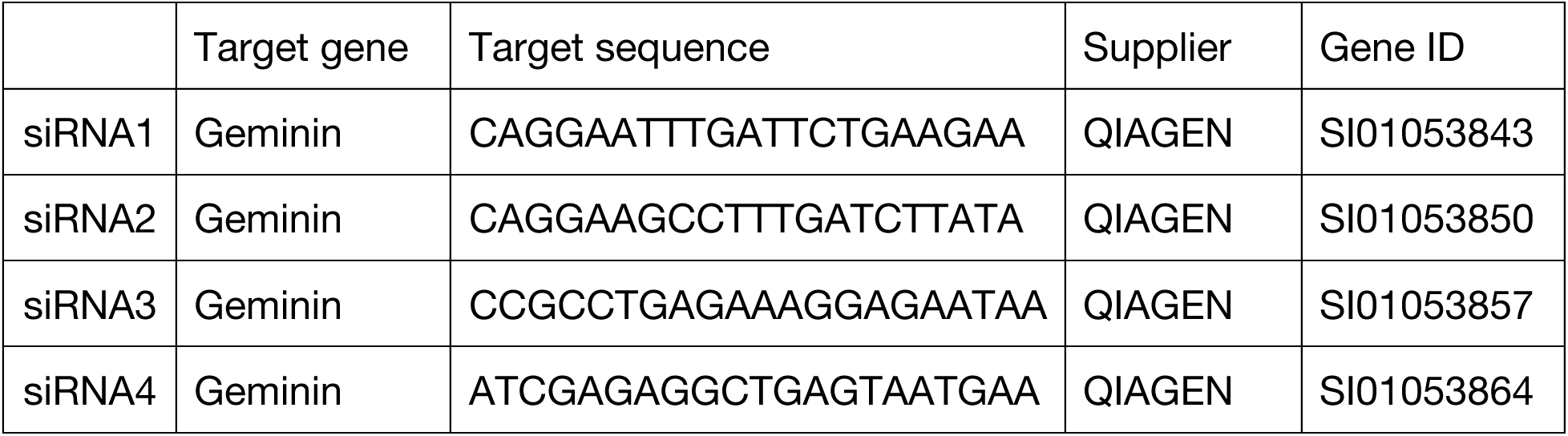
siRNAs used in this work.

**Table 2.** Inhibitors used in this work.

| Target | Inhibitor | Concentration | Supplier | Catalog No. |
|---|---|---|---|---|
| DNA–PK | KU–57788 | 1 µM | MCE | HY–11006 |
| ATM | KU–55933 | 10 µM | Selleck | S1092 |
| ATR | ETR–46464 | 0.75 µM | Selleck | S8050 |
| Chk1 | PF–477736 | 1 µM | Selleck | S2904 |
| Chk2 | BML–277 | 9 µM | Selleck | S8632 |
| Wee1 | MK–1775 | 75 nM | Selleck | S1525 |
| Plk1 | BI–2536 | 10 nM | Selleck | S1109 |
| Plk4 | LCR–263 | 150 nM | Selleck | S7837 |

### Western–blot

Cells were harvested and lysed in RIPA buffer containing 1 U/ml Pierce™ Universal Nuclease (Thermo Scientific #88700) and protease inhibitors (Roche #04693116001) and incubated on ice for 1 hour. The supernatant fraction was mixed with 4x Laemmli buffer (Bio–Rad #1610747). Equal protein amounts were subjected to electrophoresis on 15% Tris–Glycine Acrylamide gels, transferred to nitrocellulose membranes and immunoblotted with the indicated primary and secondary antibodies. Blots were developed with enhanced chemiluminescence (BioRad #1705061). Primary antibodies are listed in *Table 3*.

**Table 3.** Antibodies used in this work.

| Antibody | Raised in | Supplier | Reference |
|---|---|---|---|
| Arl13B | Rabbit | Proteintech | 17711–1–AP |
| Cep152 | Rabbit | Abcam | ab183911 |
| Cep164 | Rabbit | Proteintech | 22227–1–AP |
| Chk1 (p–Ser317) | Rabbit | Cell Signaling Technology | 12302 |
| Chk2 (p–Thr68) | Rabbit | Cell Signaling Technology | 2661 |
| GAPDH | Rabbit | Abcam | ab9485 |
| Geminin | Rabbit | Proteintech | 10802–1–AP |
| GTU88 (**γ**–tubulin) | Mouse | Sigma | T6557 |
| α–tubulin | Guinea pig | ABCD antibodies | AA345 |
| acetylated tubulin | Mouse | Sigma | T7451 |
| **γ**–H2AX (p–Ser139) | Rabbit | Abcam | ab81299 |
| Alexa Fluor™ 488 anti-rabbit | Donkey | Invitrogen | A-21206 |
| Alexa Fluor™ 568 anti-mouse | Goat | Invitrogen | A-11031 |
| Alexa Fluor™ 647 anti-rabbit | Goat | Invitrogen | A-21450 |
| HRP-conjugated anti-rabbit IgG | Goat | Jackson ImmunoResearch | 111-035-008 |

### RT–qPCR

RNA samples were prepared using the RNeasy Kit (QIAGEN 74004) according to the manufacturer’s instructions. RT–qPCR was performed using 50 ng of RNA with the Luna Universal One–Step RT–qPCR Kit (NEB E3005) on an Applied Biosystems StepOnePlus Real–Time PCR System. Each reaction was run with three technical replicates per sample. Relative gene expression was calculated using the ΔΔCt method, with Gapdh as the reference gene. Primer sequences are listed in *Table 4*.

**Table 4.**
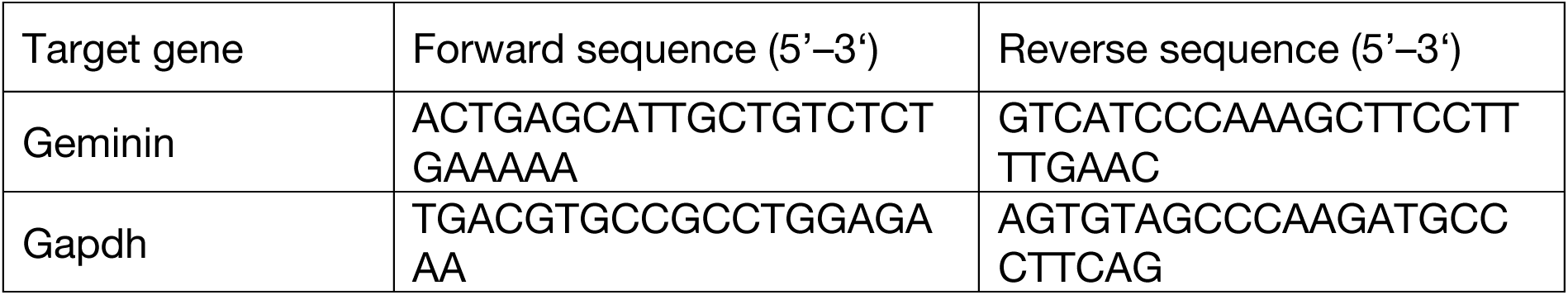
RT–qPCR primers used in this work.

### Fluorescent–activated cell sorting (FACS)

FACS analysis for DNA content was performed using standard procedures and 2 µg/mL DAPI staining. For EdU staining, cells were pulsed with 10 µM EdU for 2 hours, fixed, and processed according to the manufacturer’s protocol (Invitrogen C10424). Flow cytometry was performed using the Cytoflex S Analyzer (Beckman Coulter) and data were analyzed using CytExpert analysis software. To calculate the percentage of G0/G1, S (EdU+), and G2/M phases, and cells with DNA content above 4n, cell doublets and sub–G1 apoptotic cells were excluded.

### Immunofluorescence

Cells were grown on ⌀ 12 mm coverslips, washed with pre– warmed PBS for 3 minutes, and then fixed in ice–cold methanol for 12 minutes at –20 °C, or fixed with 4% formaldehyde for 10 minutes at room temperature. Cells were washed three times, permeabilized in PBS/0.5% Triton X–100 for 15 minutes followed by 3 washes. Blocking was performed in PBS/0.1% Tween–20/10% FBS for 30 minutes at room temperature, followed by overnight incubation with primary antibodies diluted in PBS/0.1% Tween–20/1% FBS, at 4 °C. Cells were washed three times before overnight incubation with secondary antibodies diluted in PBS/0.1% Tween–20/1% FBS, at 4 °C. DNA staining was performed with 0.5 µg/mL DAPI and cells were washed three times before mounting in Vectashield mounting medium (Vector Labs H–1900–2). Primary and secondary antibodies are listed in *Table 3*.

### Confocal microscopy

Slides were imaged on a Leica SP8 Stellaris inverted microscope with 63×/1.4 N.A. oil immersion objective using the Leica Application Suite X software (LAS–X). Images were deconvolved with Huygens Professional version 18.10 (Scientific Volume Imaging, The Netherlands). Images were processed and analyzed with ImageJ version 1.54p. Figures were created using the QuickFigures2 plugin for FIJI/ImageJ [26]. All images shown are the maximum intensity projections of all Z optical sections.

### Ultra–expansion microscopy (U–ExM)

U–ExM was performed as previously described in [25], with minor modifications. Briefly, cells were fixed with methanol as indicated above for immunofluorescence. After PBS washes, coverslips were infused with 2% acrylamide and 1.4% formaldehyde in PBS for 5 h at 37 °C. Samples were embedded in U–ExM monomer solution and gelation proceeded for 1 h at 37 °C in a humid chamber, followed by denaturation at 95 °C for 90 minutes and expansion in deionized water. Gels were immunostained with the indicated primary and secondary antibodies in PBS–based blocking buffer containing 10% FBS and 0.1% Tween–20, overnight at 4 °C. Following final expansion in deionized water, gels were mounted cell–side down on glass–bottom dishes for imaging. The protocol used achieves an expansion factor of approximately fourfold.

### Stimulated emission depletion (STED) nanoscopy

U–ExM samples were imaged on an Abberior STEDYCON 2D 775 nm (pulsed) STED assembled onto a Zeiss Observer.Z1 inverted microscope with a 100×/1.46 N.A. oil immersion objective using the STEDYCON software. Images were processed and analyzed with ImageJ version 1.54p. Figures were created using the QuickFigures2 plugin for FIJI/ImageJ [26]. All images shown are the maximum intensity projections of all Z optical sections.

### Quantification and statistical analysis

A minimum of three biological replicates were performed for each experiment. For centriole quantification, each vertical bar in the graph represents the average of all the cells at that stage from a single field of view, typically 20–30 cells, with at least 200 cells quantified, per condition. Data were analyzed using nonparametric statistical methods. Differences across multiple groups were evaluated with the Kruskal–Wallis test followed by Dunn’s multiple comparisons test for multiple pairwise comparisons. For nuclear foci quantification, ImageJ was used to segment isolated nuclei and quantify the number of foci within each. Nuclei were then scored for positive or negative EdU incorporation, to distinguish S phase cells. Statistical tests are indicated in figure legends. For centriolar distances measurements, individual measurements from three independent biological replicates were pooled for each experimental condition. Data distributions were assessed using the Shapiro–Wilk test and by comparing Gaussian and lognormal distributions. Data were log10–transformed prior to statistical analysis, and homogeneity of variance was assessed using the Brown–Forsythe test. Log10–transformed data were compared using one–way ordinary ANOVA followed by Šídák’s multiple–comparisons test. Statistical analyses were performed on transformed measurements, while untransformed normalized values were used for graphical representation. Distances were normalized to the mean control distance and are presented as fold change relative to control. All data were plotted and analyzed using GraphPad Prism 11.0.2.

## Data Availability

This study includes no data deposited in external repositories. The data supporting the findings of this study are available within the article and its supplementary information.

## Author contributions

Conceptualization, IBS, DMG; Investigation, IBS; Analysis, IBS; Writing, Review & Editing, IBS, DMG; Funding acquisition, DMG.

## Acknowledgments

We are grateful to Paula Almeida Coelho for helpful discussions, and to Vickie Yang for assistance with the U–ExM method used in this study. We thank Caltech Biological

Imaging Facility and Caltech Flow Cytometry and Cell Sorting Facility for resource support. Research reported in this publication was supported by the National Cancer Institute of the National Institutes of Health under award number 5R01CA259382–06.

---

**Supplementary Figure 1.**
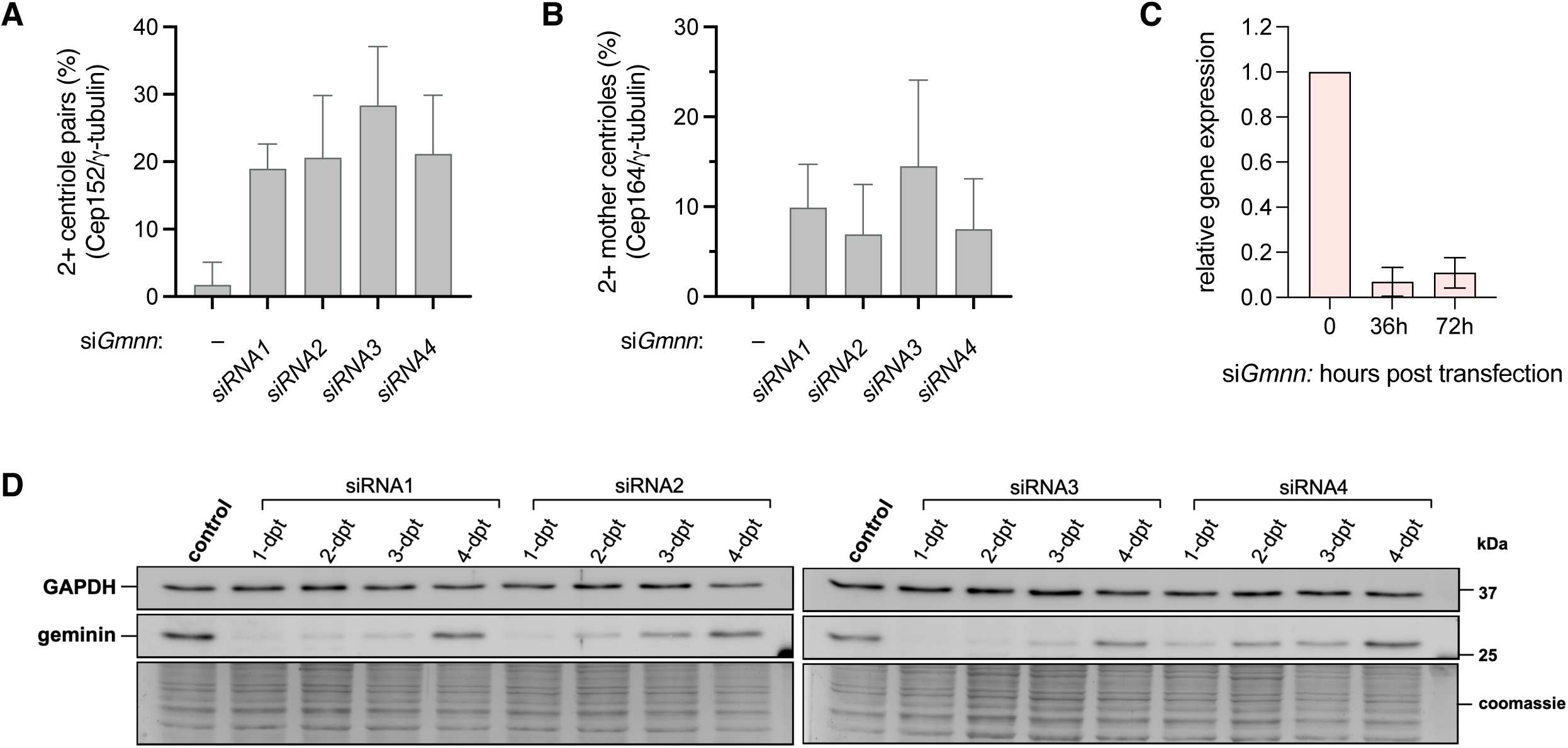
Validation of Geminin–depletion and individual siRNA phenotypes. (A) Graphic representation of the percentage of cells showing more than 2 (>2) centriole pairs, per cell (centrosome amplification phenotype). MEFs were stained to reveal Cep152, **γ**–tubulin, *a*–tubulin and DNA. Colocalization of Cep152 with **γ**–tubulin identifies centriole pairs/centrosomes. 4 siRNA oligonucleotides against Geminin were tested for their efficacy to induce centriole/centrosome amplification in MEFs. (B) Graphic representation of the percentage of cells showing more than 2 (>2) mother centrioles, per cell (centrosome amplification phenotype). MEFs are stained to reveal Cep164, **γ**–tubulin, *a*–tubulin and DNA. Colocalization of Cep164 with **γ**–tubulin identifies mother centrioles. 4 siRNA oligonucleotides against Geminin were tested for their efficacy to induce mother centriole amplification in MEFs. (C) RT–qPCR analysis of relative levels of Geminin mRNA transcripts in MEFs treated with siRNA against Geminin at the indicated timepoints. The average and standard deviation (SD) from two biological samples, three technical replicates within each, are plotted. (D) Western–blot analysis of Geminin protein levels in MEFs treated with 4 siRNA oligonucleotides against Geminin. siRNA1–4 were individually tested for their efficacy to deplete endogenous Geminin up to 4 dpt.

**Supplementary Figure 2.**
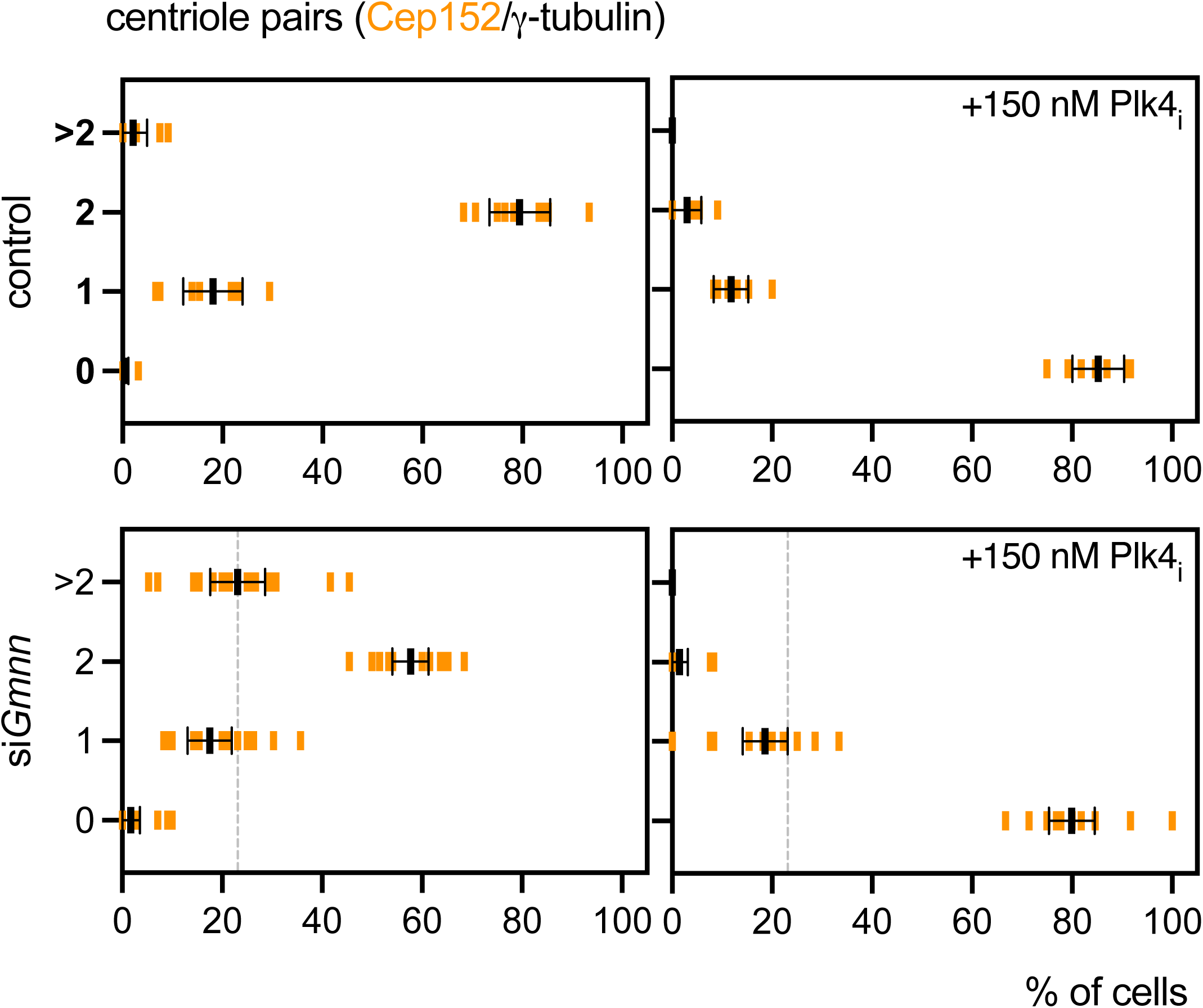
Plk4 inhibitor validates centriole quantification assays. Graphic representation of the percentage of cells showing 0, 1, 2 and more than 2 (>2) centriole pairs, per cell. Control and Geminin–depleted MEFs were stained to reveal Cep152, **γ**–tubulin, *a*–tubulin and DNA. Colocalization of Cep152 with **γ**–tubulin identifies centriole pairs/centrosomes. MEFs were treated with the Plk4 inhibitor, centrinone/LCR–263. The number of centrioles was quantified for each category, and percentage values are shown as vertical bars. Experiments were performed at least 3 times independently. Error bars correspond to mean ± 95% confidence interval (CI).Plk4_i_ was added to cells in culture at 1 dpt (inhibitor treatment time: 3 days prior to fixation).

## References

[1] Tsou MF, Wang WJ, George KA, Uryu K, Stearns T, Jallepalli PV. Polo kinase and separase regulate the mitotic licensing of centriole duplication in human cells. Dev Cell. 2009 Sep;17(3):344–54. doi: 10.1016/j.devcel.2009.07.015. PMID: 19758559; PMCID: PMC2746921.

[2] Tsou MF, Stearns T. Mechanism limiting centrosome duplication to once per cell cycle. Nature. 2006 Aug 24;442(7105):947–51. doi: 10.1038/nature04985. Epub 2006 Jul 19. PMID: 16862117.

[3] Kim J, Lee K, Rhee K. PLK1 regulation of PCNT cleavage ensures fidelity of centriole separation during mitotic exit. Nat Commun. 2015 Dec 9;6:10076. doi: 10.1038/ncomms10076. PMID: 26647647; PMCID: PMC4682042.

[4] Stillman B, Diffley JFX, Iwasa JH. Mechanisms for licensing origins of DNA replication in eukaryotic cells. Nat Struct Mol Biol. 2025 Jul;32(7):1143–1153. doi: 10.1038/s41594-025-01587-5. Epub 2025 Jun 30. PMID: 40588663; PMCID: PMC12335275.

[5] Wohlschlegel JA, Dwyer BT, Dhar SK, Cvetic C, Walter JC, Dutta A. Inhibition of eukaryotic DNA replication by geminin binding to Cdt1. Science. 2000 Dec 22;290(5500):2309–12. doi: 10.1126/science.290.5500.2309. PMID: 11125146.

[6] Tomkins J, Edwardes LV, Faull SV, Peach M, Gillespie PJ, Leber V, Schmidt A, Bounoua H, Sim N, Camarillo R, Blow JJ, Barr AR, Barnard A, Speck C. Geminin inhibits DNA replication licensing by sterically blocking CDT1-MCM2 interactions. Nat Commun. 2025 Dec 9;16(1):11040. doi: 10.1038/s41467-025-67073-0. PMID: 41365879; PMCID: PMC12695944.

[7] McGarry TJ, Kirschner MW. Geminin, an inhibitor of DNA replication, is degraded during mitosis. Cell. 1998 Jun 12;93(6):1043–53. doi: 10.1016/s0092-8674(00)81209-x. PMID: 9635433.

[8] Gorgoulis VG, Vassiliou LV, Karakaidos P, Zacharatos P, Kotsinas A, Liloglou T, Venere M, Ditullio RA Jr, Kastrinakis NG, Levy B, Kletsas D, Yoneta A, Herlyn M, Kittas C, Halazonetis TD. Activation of the DNA damage checkpoint and genomic instability in human precancerous lesions. Nature. 2005 Apr 14;434(7035):907–13. doi: 10.1038/nature03485. PMID: 15829965.

[9] Godinho SA, Pellman D. Causes and consequences of centrosome abnormalities in cancer. Philos Trans R Soc Lond B Biol Sci. 2014 Sep 5;369(1650):20130467. doi: 10.1098/rstb.2013.0467. PMID: 25047621; PMCID: PMC4113111.

[10] Burrage LC, Charng WL, Eldomery MK, Willer JR, Davis EE, Lugtenberg D, Zhu W, Leduc MS, Akdemir ZC, Azamian M, Zapata G, Hernandez PP, Schoots J, de Munnik SA, Roepman R, Pearring JN, Jhangiani S, Katsanis N, Vissers LE, Brunner HG, Beaudet AL, Rosenfeld JA, Muzny DM, Gibbs RA, Eng CM, Xia F, Lalani SR, Lupski JR, Bongers EM, Yang Y. De Novo GMNN Mutations Cause Autosomal–Dominant Primordial Dwarfism Associated with Meier–Gorlin Syndrome. Am J Hum Genet. 2015 Dec 3;97(6):904–13. doi: 10.1016/j.ajhg.2015.11.006. PMID: 26637980; PMCID: PMC4678788.

[11] Nielsen-Dandoroff E, Ruegg MSG, Bicknell LS. The expanding genetic and clinical landscape associated with Meier–Gorlin syndrome. Eur J Hum Genet. 2023 Aug;31(8):859–868. doi: 10.1038/s41431-023-01359-z. Epub 2023 Apr 14. PMID: 37059840; PMCID: PMC10400559.

[12] Klotz-Noack K, McIntosh D, Schurch N, Pratt N, Blow JJ. Re-replication induced by geminin depletion occurs from G2 and is enhanced by checkpoint activation. J Cell Sci. 2012 May 15;125(Pt 10):2436–45. doi: 10.1242/jcs.100883. Epub 2012 Feb 24. PMID: 22366459; PMCID: PMC3481538.

[13] Saldivar JC, Hamperl S, Bocek MJ, Chung M, Bass TE, Cisneros-Soberanis F, Samejima K, Xie L, Paulson JR, Earnshaw WC, Cortez D, Meyer T, Cimprich KA. An intrinsic S/G_2_checkpoint enforced by ATR. Science. 2018 Aug 24;361(6404):806–810. doi: 10.1126/science.aap9346. PMID: 30139873; PMCID: PMC6365305.

[14] Michelena J, Gatti M, Teloni F, Imhof R, Altmeyer M. Basal CHK1 activity safeguards its stability to maintain intrinsic S–phase checkpoint functions. J Cell Biol. 2019 Sep 2;218(9):2865–2875. doi: 10.1083/jcb.201902085. Epub 2019 Jul 31. PMID: 31366665; PMCID: PMC6719454.

[15] Mihaylov IS, Kondo T, Jones L, Ryzhikov S, Tanaka J, Zheng J, Higa LA, Minamino N, Cooley L, Zhang H. Control of DNA replication and chromosome ploidy by geminin and cyclin A. Mol Cell Biol. 2002 Mar;22(6):1868–80. doi: 10.1128/MCB.22.6.1868-1880.2002. PMID: 11865064; PMCID: PMC135598.

[16] Melixetian M, Ballabeni A, Masiero L, Gasparini P, Zamponi R, Bartek J, Lukas J, Helin K. Loss of Geminin induces rereplication in the presence of functional p53. J Cell Biol. 2004 May 24;165(4):473–82. doi: 10.1083/jcb.200403106. PMID: 15159417; PMCID: PMC2172361.

[17] Harvey SL, Charlet A, Haas W, Gygi SP, Kellogg DR. Cdk1–dependent regulation of the mitotic inhibitor Wee1. Cell. 2005 Aug 12;122(3):407–20. doi: 10.1016/j.cell.2005.05.029. PMID: 16096060.

[18] Dwivedi D, Harry D, Meraldi P. Mild replication stress causes premature centriole disengagement via a sub–critical Plk1 activity under the control of ATR–Chk1. Nat Commun. 2023 Sep 29;14(1):6088. doi: 10.1038/s41467-023-41753-1. PMID: 37773176; PMCID: PMC10541884.

[19] Lemmens B, Hegarat N, Akopyan K, Sala-Gaston J, Bartek J, Hochegger H, Lindqvist A. DNA Replication Determines Timing of Mitosis by Restricting CDK1 and PLK1 Activation. Mol Cell. 2018 Jul 5;71(1):117–128.e3. doi: 10.1016/j.molcel.2018.05.026. Epub 2018 Jun 28. PMID: 30008317; PMCID: PMC6039720.

[20] Berti M, Cortez D, Lopes M. The plasticity of DNA replication forks in response to clinically relevant genotoxic stress. Nat Rev Mol Cell Biol. 2020 Oct;21(10):633–651. doi: 10.1038/s41580-020-0257-5. Epub 2020 Jul 1. PMID: 32612242.

[21] Xu X, Weaver Z, Linke SP, Li C, Gotay J, Wang XW, Harris CC, Ried T, Deng CX. Centrosome amplification and a defective G2-M cell cycle checkpoint induce genetic instability in BRCA1 exon 11 isoform-deficient cells. Mol Cell. 1999 Mar;3(3):389–95. doi: 10.1016/s1097-2765(00)80466-9. PMID: 10198641.

[22] Mardin BR, Agircan FG, Lange C, Schiebel E. Plk1 controls the Nek2A-PP1γ antagonism in centrosome disjunction. Curr Biol. 2011 Jul 12;21(13):1145–51. doi: 10.1016/j.cub.2011.05.047. Epub 2011 Jun 30. PMID: 21723128.

[23] Gaillard H, García-Muse T, Aguilera A. Replication stress and cancer. Nat Rev Cancer. 2015 May;15(5):276–89. doi: 10.1038/nrc3916. PMID: 25907220.

[24] da Costa AABA, Chowdhury D, Shapiro GI, D’Andrea AD, Konstantinopoulos PA. Targeting replication stress in cancer therapy. Nat Rev Drug Discov. 2023 Jan;22(1):38–58. doi: 10.1038/s41573-022-00558-5. Epub 2022 Oct 6. PMID: 36202931; PMCID: PMC11132912.

[25] Coelho PA, Fatalska A, Geymonat M, Lattao R, Glover DM. Sensing centrosome amplification: the interface between centriole duplication and autophagy. Nat Commun. 2026 Jun 21. doi: 10.1038/s41467-026-74702-9. Epub ahead of print. PMID: 42324259.

[26] Mazo G. QuickFigures: A toolkit and ImageJ PlugIn to quickly transform microscope images into scientific figures. PLoS One. 2021 Nov 9;16(11):e0240280. doi: 10.1371/journal.pone.0240280. PMID: 34752463; PMCID: PMC8577749.

[27] Gambarotto D, Zwettler FU, Le Guennec M, Schmidt-Cernohorska M, Fortun D, Borgers S, Heine J, Schloetel JG, Reuss M, Unser M, Boyden ES, Sauer M, Hamel V, Guichard P. Imaging cellular ultrastructures using expansion microscopy (U– ExM). Nat Methods. 2019 Jan;16(1):71–74. doi: 10.1038/s41592-018-0238-1. Epub 2018 Dec 17. PMID: 30559430; PMCID: PMC6314451.

